# Agarose Stamped Device: A simple but effective fixation method for Zebrafish Larvae through customizable 3D printed molds for *Danio rerio* (Zebrafish)

**DOI:** 10.1101/2025.03.04.641502

**Authors:** John Jutoy, Hossein Mehrabi, Erica Jung

## Abstract

Precise immobilization of zebrafish larvae is critical for high-resolution neural imaging and behavioral studies, yet conventional methods are often labor-intensive, low-throughput, and can induce stress. To address these challenges, we introduce the Agarose Stamped Device (ASD), a novel platform enabling efficient and gentle immobilization of zebrafish larvae for high-resolution imaging and behavioral assays. By employing a stamping approach to mold an array of consistent agarose wells, the ASD enables rapid, parallel, and reproducible positioning of larvae in a standardized orientation while minimizing stress and preserving viability. We validated ASD performance by comparing larval survival, heart rate, and imaging stability to conventional agarose embedding, finding no adverse effects on larval physiology and significantly enhanced imaging throughput and data consistency. Demonstrations in neuroscience and behavioral experiments underscore the device’s versatility for diverse in vivo studies. The ASD offers a simple, cost-effective, and powerful tool for advancing zebrafish-based neuroscience and behavioral research.

## Introduction

Larval zebrafish (Danio rerio) have become an indispensable model for neuroscience and behavioral research due to their genetic tractability, small size, and optical transparency^1–3^. Foundational studies established that zebrafish larvae can model complex biological processes from neurodevelopment to behavior, enabling high-throughput drug screens and neurobiological assays in a whole-organism context^2,4^. Early behavioral assays often employed free-swimming larvae in multi-well plates or Petri dishes to observe locomotor responses and drug effects.

However, while these open-well methods are straightforward, they offer limited experimental control – stimulus delivery and behavioral readouts tend to be qualitative and inconsistent^5^. As the field has advanced, there is a growing need for methods to restrain or position larvae in reproducible orientations for high-resolution imaging and precise stimulus presentation, without compromising the animal’s viability or normal behavior.

The conventional approach for immobilizing zebrafish larvae is to embed them in low-melting-point agarose. This simple, low-barrier method provides adequate immobilization for microscopy or targeted stimulation and has been widely used in neurobehavioral experiments. Unfortunately, agarose embedding presents several well-recognized challenges. Fully encasing a larva in agarose can impede its growth or physiological processes and has been reported to induce stress (e.g. cardiac strain, tissue necrosis) when maintained for prolonged periods^6,7^. Standard agarose mounts also solidify rapidly, giving researchers only a brief window to orient each larva under the microscope. Manipulating larvae into position before the gel sets requires considerable skill and risks physical damage to the specimen^8,9^. Moreover, once solidified, the agarose may need to be partially removed to allow certain behaviors or growth to proceed, a process that itself can injure the larva^8^. These drawbacks limit the utility of plain agarose for long-term behavioral studies or experiments requiring frequent interventions.

To overcome such issues, researchers have explored both low-barrier and high-barrier entry methods for zebrafish larval immobilization. On the low-barrier end, simple innovations like casting agarose wells or “divots” in advance have been used to guide larval positioning^6^. For example, ridged agarose molds can hold larvae in arrayed rows, reducing the hands-on alignment needed before imaging^6^. Such approaches retain the low cost and accessibility of agarose while improving consistency. However, they still involve manual preparation and do not completely resolve limitations in long-term viability or access to the specimen. On the high-barrier end, advanced microfluidic and engineered devices have recently been developed to precisely position and hold zebrafish larvae. Notably, Beebe and colleagues introduced the ZEBRA (Zebrafish Entrapment by Restriction Array) microfluidic device as an agarose-free mounting technique that uses only a pipette for loading larvae into preset channels^10^. This and other “fish-on-a-chip” systems provide fine spatial control, allowing parallel orientation of multiple larvae and easy delivery of stimuli or reagents via built-in ports^10^. Such high-end devices have enabled new experimental paradigms – for instance, microfluidic platforms can expose larvae to controlled flow or electric fields to study rheotaxis and electrotaxis behaviors with quantitative precision^5^. They have also been applied to high-throughput drug screens and whole-brain activity mapping in vivo^11^. The downside is that these setups often require specialized fabrication and can pose a higher barrier to entry for many labs. Indeed, one persistent challenge with microfluidic larval holders is limited throughput; handling and imaging larvae one-by-one (or only a few at a time) can be time-consuming compared to traditional multi-well assays^12^. Recent advancements are beginning to address this by scaling up device capacity (e.g. from single-larva to four-larva chips) and streamlining loading procedures^12^, but the complexity and cost of microdevices still motivate the search for simpler alternatives.

In this context, we introduce the Agarose Stamped Device (ASD) as a novel solution for zebrafish larval alignment and fixation. The ASD bridges the gap between low-barrier agarose embedding and higher-barrier microfluidic devices. It consists of a patterned stamp that imprints an array of shallow larva-sized wells into agarose, creating a ready-to-use template for positioning larvae. Using the ASD, a researcher can immobilize dozens of larvae in one step by simply placing them into the stamped wells, where the animals naturally settle into the correct orientation^13^. This approach offers several key advantages over existing methods. First, it drastically reduces handling time and user skill required – there is no need to individually position each larva, as they self-orient within the molded wells^13^. Second, because each well is identical, the larvae are uniformly aligned (for example, all facing the same direction and lying on the same focal plane), which improves imaging consistency and assay reproducibility^13^.

Third, the stamped agarose substrate leaves the larvae partly exposed to the surrounding solution, creating an open system that permits easy access for experimental manipulations^13^. For instance, drugs or stimuli can be added to the medium at any time, and specific body parts (e.g. the tail or eyes) can be targeted for stimulation or recording, something not possible when a larva is fully embedded in agarose. Importantly, the gentle physical restraint provided by the ASD is sufficient to hold larvae in place for microscopy or behavioral monitoring, while minimizing stress and allowing normal physiological functions (heartbeat, blood flow, neural activity) to continue unimpeded. Previous implementations of agarose stamping have demonstrated that larvae up to 20 days post-fertilization can be stably maintained and imaged with such devices^13^, underscoring the method’s applicability across developmental stages.

By combining low technical barriers with high positioning precision, the ASD opens new opportunities for zebrafish behavioral experiments. It enables high-content analyses where many larvae are immobilized in parallel – for example, drug screening assays can treat multiple aligned larvae with compounds and directly compare their responses under the same optical field^13^. Likewise, neurobehavioral studies requiring simultaneous imaging of neural activity in several animals (such as brain calcium imaging during sensory stimulation) can benefit from the standardized orientation and stability the ASD provides. In this work, we validate the ASD’s effectiveness in reliably fixing larvae in place and maintaining consistent alignment across individuals. We demonstrate that larvae mounted with the ASD remain properly aligned and exhibit expected behaviors or responses, indicating that the device imposes minimal adverse effects. Overall, the Agarose Stamped Device offers a convenient and reproducible immobilization strategy that addresses many challenges of conventional agarose embedding. We propose that the ASD will facilitate a wide range of zebrafish larval assays – from developmental imaging to behavioral neuroscience – by providing an accessible yet robust means of larval fixation and alignment. The following sections detail the design of the ASD, its performance relative to established methods, and example applications that highlight its utility in zebrafish behavioral research.

## Results and Discussion

### Agarose Stamping Platform Workflow Fabrication of Stamp

The stamp was resin 3D printed with an Elegoo Mars 2 Pro. It was then submerged in a 75% isopropyl alcohol bath to remove any excess uncured resin. After further drying, the stamp was further cured with an ultraviolet light (670 mm wavelength) for one minute on each side to solidify the stamp.

### Creation of Agarose Stamp Device

A 1.5% Agarose solution was made with Sigma Aldrich. The solution was poured into a base plate of a 35 mm petri dish containing the stamp such that the base of the stamp was facing up. The Agarose device was left to solidify for 15 minutes, from which the stamp was carefully removed. For best results, a small knife was used to cut any Agarose in contact with the stamps base, and the stamp was slowly lifted with a pair of tweezers.

### Hardware: Agarose Stamping Overview

The fabrication of agarose stamping devices involves several key steps. The first step is the design process using CAD software (Figure 1A). Based on experimental requirements, essential features are selected to optimize the device’s functionality. These features may include: single stamps for experiments involving individual larvae, array stamps for high-throughput experiments with multiple larvae (e.g., screening), microchannels to interconnect the slots for fluidic experiments, reservoirs for long-term studies requiring stable water levels, and a free-tail configuration for studies where tail movement must be recorded (Figure 1A.1). After selecting the necessary features, the design is detailed in CAD software (Figure 1A.2), and the final model is exported as an STL file for mold fabrication (Figure 1B).

**Figure 1.**
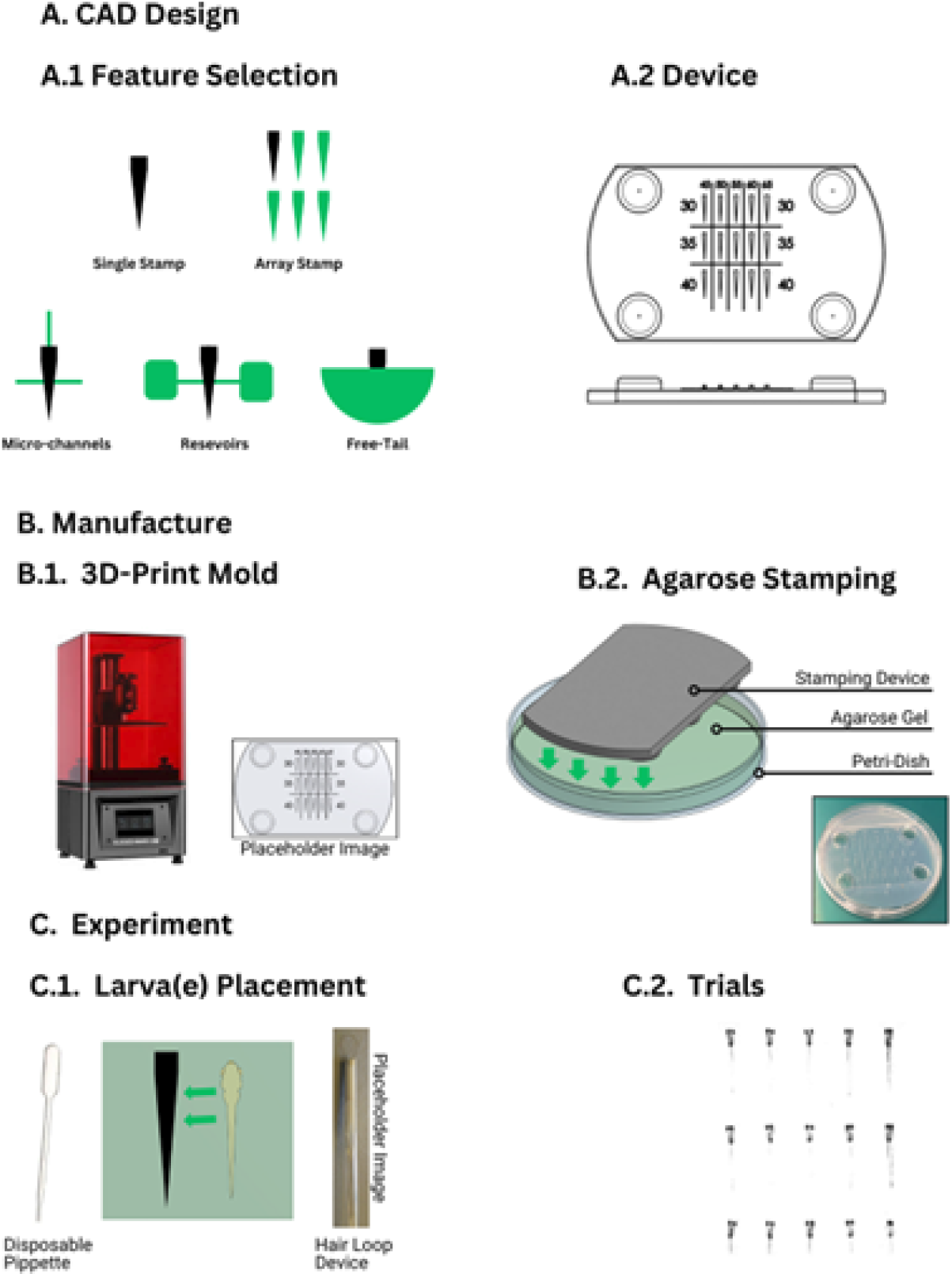
Agarose Stamping Workflow. The Agarose Stamping Workflow visualized into three sections: (A) CAD Design, (B) Device Fabrication, and (C) Device Utilization. (A.1) Various features can augment to the single stamp setup to accommodate requirements for experiments requiring high throughput, fluid channels, fluid reservoirs, and free tail configurations. A high throughput device is displayed in the figure. (A.2) Once features have been chosen, they can be integrated together through CAD software to create a stamp model. Note that four extruded circles were included to act as a spacer from the agarose (B.1) The model can be printed with a resin printer that has at least 50 microns of resolution. (B.2) The printed stamp is placed on a petri dish and is submerged up to the stamp’s base with heated liquid agarose. (C.1) Larva is placed near the hole (negative larva shape that was made with the stamp) via disposable pipette. The larva is maneuvered into the hole by moving it with the loop of a hair loop device ensuring that the head of the larvae is aligned with the hole. (C.2) High throughput device ran through a visual stimulation trial.

**Figure 2.**
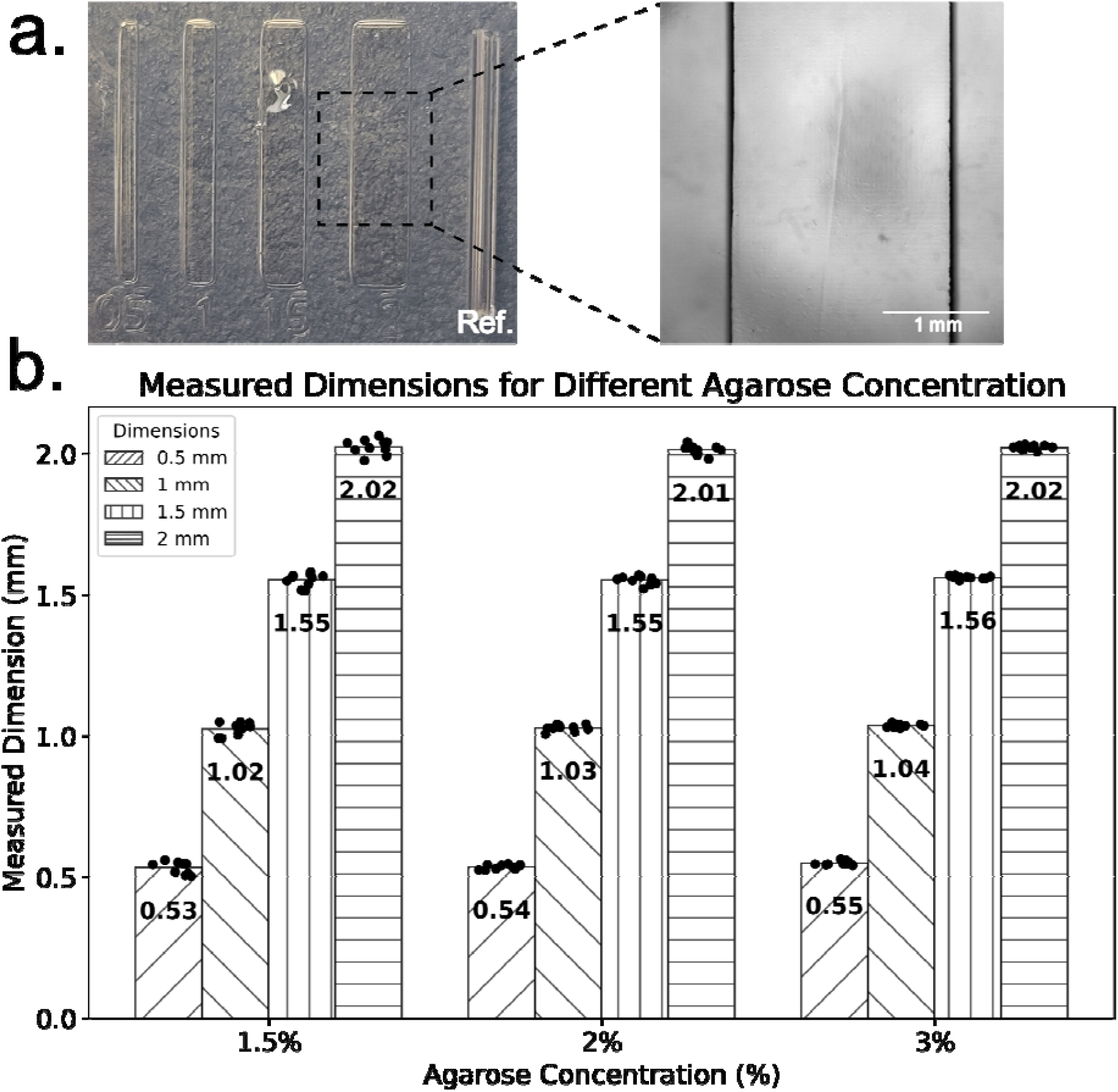
Evaluation of feature dimension fidelity in agarose stamps. (a) Microfabricated features with nominal widths of 0.5, 1, 1.5, and 2 mm imprinted in agarose using a 3D-printed mold. A 0.5 mm reference feature is included in each set for dimensional calibration. A magnified view of the 2 mm feature is shown, with a scale bar of 1 mm for reference. (b) Measured feature dimensions for agarose stamps fabricated with three different concentrations (1.5%, 2%, and 3%). Each bar represents the mean measured dimension (N = 10) for each nominal feature width (0.5, 1, 1.5, and 2 mm), with individual data points overlaid. The numerical values on each bar indicate the mean feature dimension.

**Figure 3.**
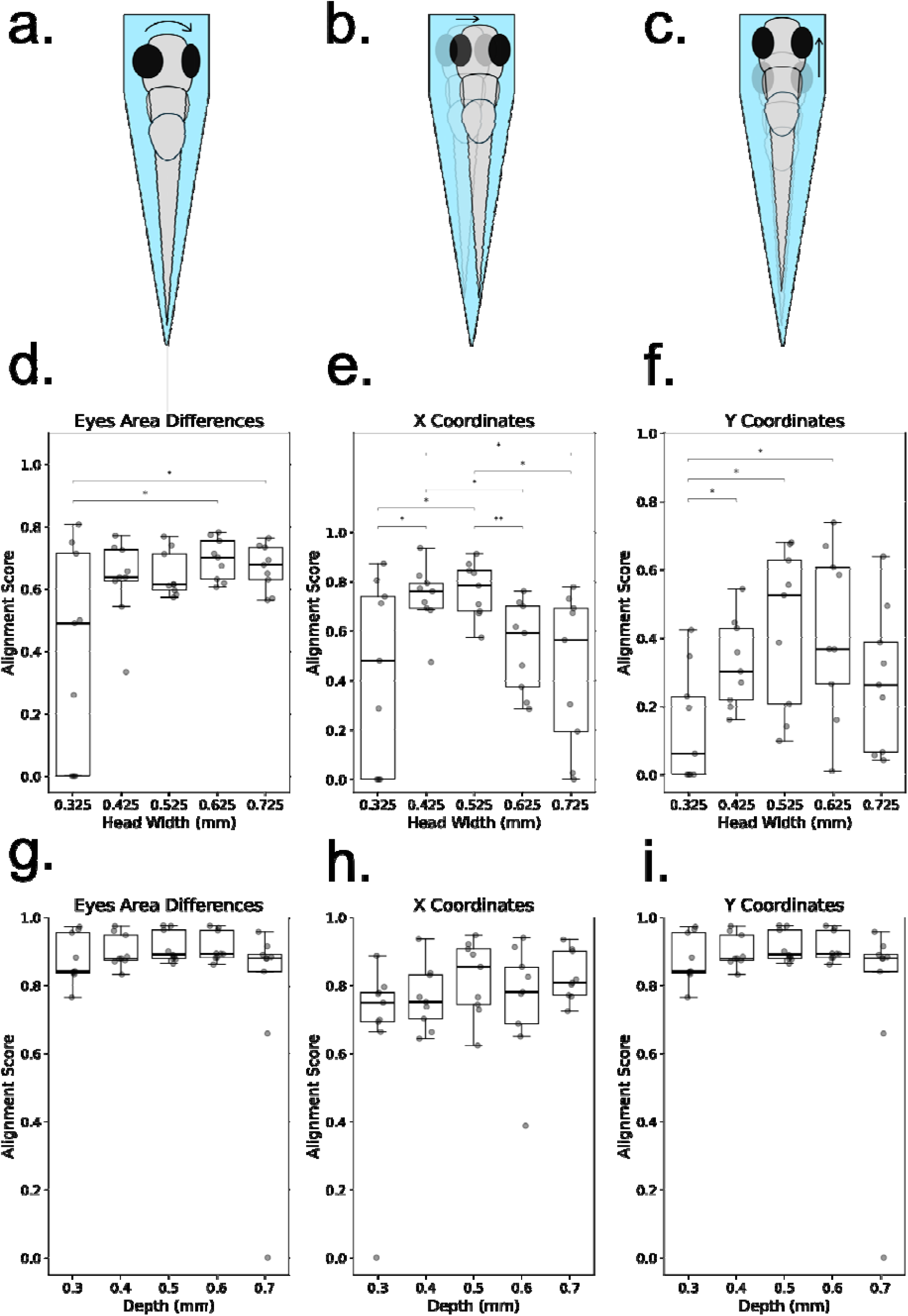
Influence of zebrafish larval positioning on alignment metrics in a microfluidic channel. (a–c) Schematic representations of common misalignments and instabilities observed when zebrafish larvae are positioned in a microfluidic channel: (a) **Rotation along the anteroposterior (AP) axis**, identifiable by asymmetrical eye positioning; (b) **Mediolateral (ML) displacement**, characterized by lateral shifts in eye positions along the x-axis; (c) **Anteroposterior (AP) displacement**, observed as shifts in eye positions along the y-axis. (d–f) Alignment scores for zebrafish larvae with different head widths at a fixed channel depth, quantified using three criteria: (d) **Eye area asymmetry**, where significant differences are observed between groups 1 and 4 (*p* = 0.02706) and groups 1 and 5 (*p* = 0.03910) (Welch’s *t*-test); (e) **X-axis displacement**, showing significant differences for multiple comparisons: group 1 vs. 2 (*p* = 0.04048), group 1 vs. 3 (*p* = 0.02865), group 2 vs. 4 (*p* = 0.01505), group 2 vs. 5 (*p* = 0.02234), group 3 vs. 4 (*p* = 0.00664), and group 3 vs. 5 (*p* = 0.01471) (Welch’s *t*-test); (f) **Y-axis displacement**, with statistically significant differences observed for groups 1 vs. 2 (*p* = 0.02648), groups 1 vs. 3 (*p* = 0.01294), and groups 1 vs. 4 (*p* = 0.01655) (Mann-Whitney *U* test). (g–i) Alignment scores for zebrafish larvae with different depths at a fixed head width, assessed using the same three criteria: (g) **Eye area asymmetry**, (h) **X-axis displacement**, and (i) **Y-axis displacement**. Individual data points are displayed on each box plot, with statistical significance indicated.

**Figure 4.**
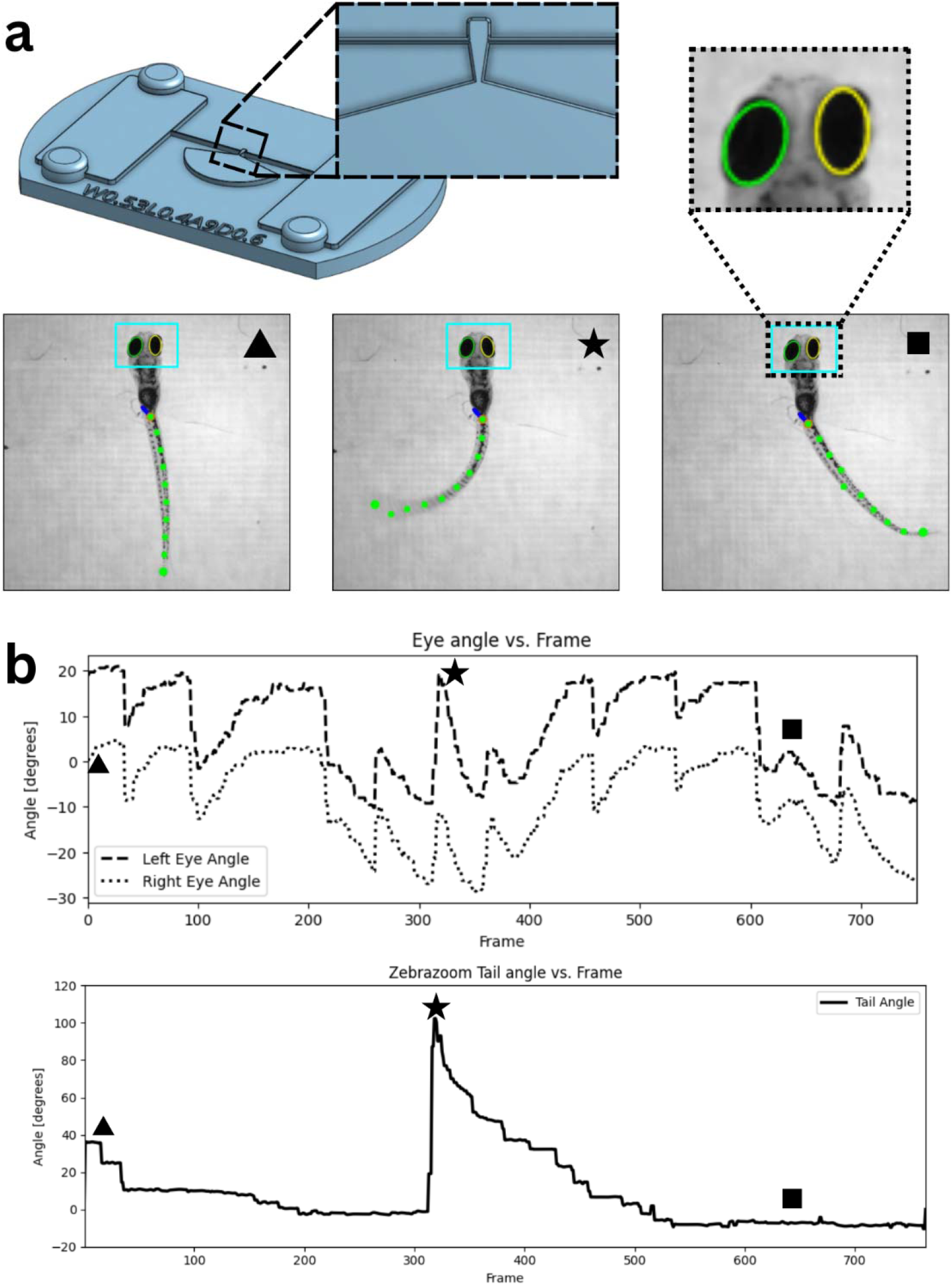
Larvae eye and tail tracking. A head fixed with free tail feature was designed to test eye and tail tracking with an optokinetic response protocol. **a** Device design and key frames from trial. A recorded video of this trial was passed through Zebrazoom for tail tracking.

The mold can be 3D printed using a variety of printers, provided the resolution is at least 50 microns (Figure 1B.1). After printing the mold, it is placed into a petri dish, and melted agarose is poured between the mold and the dish such that the agarose reaches the top edge of the mold. After the agarose solidifies, the mold is carefully removed to avoid damaging the agarose structure (Figure 1B.2). Now, the device is ready for use in experiments (Figure 1C).

For the experimental setup, larvae are transferred to their designated spots on the agarose stamp. A disposable pipette is used to initially transfer larvae near their allocated slots. Then, a hair loop tool is employed to gently position the larvae into their specific slots, ensuring they are placed without causing damage (Figure 1C.1). After all larvae are positioned, the experiment can commence (Figure 1C.2).

### Alignment Metrics

Zebrafish larvae exhibit size variations based on factors such as age, water temperature, stocking density, water quality, genetic background, and food availability^14,15^. In this study, we standardized these parameters and used 7 days post-fertilization (dpf) larvae. To perform various experiments, it is necessary to immobilize the larvae within a slot while maintaining their viability and natural behavior. The slot dimensions had to be optimized to ensure that the larvae are neither excessively constrained, which could cause damage or alter their physiological state, nor too loosely positioned, which would allow uncontrolled movement.

To determine the optimal slot dimensions, we varied both head width and slot depth. A smaller head width and increased depth resulted in a tighter fit, imposing greater restriction, while a larger head width and shallower depth provided greater freedom of movement. We examined head widths ranging from 0.325 mm to 0.725 mm in 0.1 mm increments and slot depths from 0.3 mm to 0.7 mm with the same increment. When evaluating the effect of head width, the depth was fixed at 0.5 mm, and when analyzing depth, the head width was fixed at 0.525 mm. For each head width and depth combination, nine samples were analyzed, resulting in a total of 90 samples across all conditions.

After positioning the larvae in the slot, their alignment and stability were recorded from a top-view perspective at 35 frames per second (fps) for 10 minutes. The device design incorporated water reservoirs to maintain a stable water level within the slots and microchannels, preventing dehydration or stress over time.

### Alignment and Stability Quantification

The alignment and stability of each larva were assessed based on eye positioning, as the eyes serve as clear and quantifiable markers. A well-aligned larva exhibits symmetrical eyes from the top view, and a stable larva maintains this symmetry over time. Misalignment or instability can manifest as:

1. Rotation along the anteroposterior (AP) axis, causing asymmetry in eye positioning.
2. Mediolateral (ML) displacement, resulting in lateral eye shifts along the x-axis.
3. Anteroposterior (AP) displacement, leading to movement along the y-axis.

To quantify these misalignments, we analyzed eye area differences and x/y coordinate variations over time.

### Misalignment Measurement: Eye Area Differences

When the larva rotates along the AP axis, the eyes become asymmetric, and the absolute difference in eye areas becomes nonzero. A higher value indicates greater misalignment. For each frame *i* in a sample, we calculated:

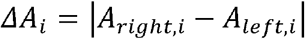

Where *A*_*right,i*_ and *A*_*left,i*_ are right and left eye area in frame *i*.

The mean eye area per sample was determined as:

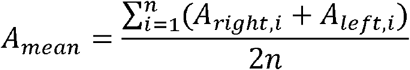

The normalized eye area difference was then computed as:

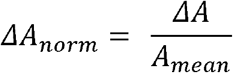

To assess variation across frames, we computed the standard deviation of the eye area differences:

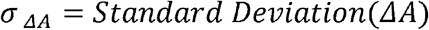

For each experimental condition, we analyzed 45 samples and determined:

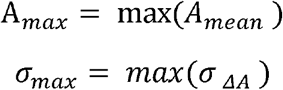

### Handling Overlapping Eyes

If the eyes overlap due to misalignment, the algorithm cannot distinguish between them, and the eye area measurement is assigned NaN. To quantify this, we calculated the NaN ratio:

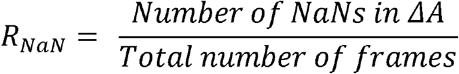

### Distinguishing Misalignment from Instability

Eye area differences may result from misalignment, which is accounted for in the normalized mean, or instability, which is accounted for in the normalized standard deviation. To separate these effects, we computed:

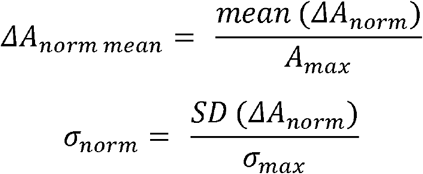

Finally, the alignment score was computed as:

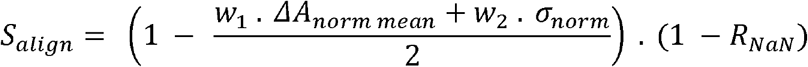

Where *w*_1_ and *w*_2_ are weights that sum to 1. We used *w*_1_= *w*_2_ =0.5, and the results remained consistent across different values.

### Instability Measurement: Eye Coordinate Variations

When the larva experiences ML or AP displacement, the x/y coordinates of the eyes fluctuate over time. The same analysis used for eye area differences was applied, with the exception that the average was not considered, only the standard deviation:

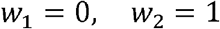

Instead of using eye area differences, we computed eye coordinate averages:

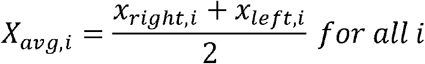

The sample mean for x-coordinates was:

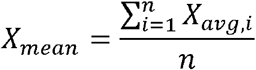

Similarly, for y-coordinates:

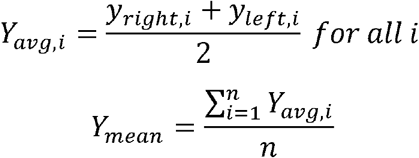

And the rest of the process follows the same methodology as the eye area differences.

### Food Particle Suction and Velocity Analysis in Zebrafish Larvae

Another application of the agarose stamping device is its ability to mechanically and physically immobilize zebrafish larvae while allowing their mouth to move freely, preserving natural feeding behavior. This feature is particularly useful for studying food intake dynamics in zebrafish larvae under controlled conditions. By trapping a 6-day post-fertilization (6 dpf) ABWT larva in a microfluidic slot connected to a microchannel (Figure 5a, b), we introduced a 0.2 mL solution of food particles into the system. Initially, the food particles remained suspended in water, and whenever the larva initiated suction, the particles exhibited a velocity proportional to the suction frequency and strength.

**Figure 5.**
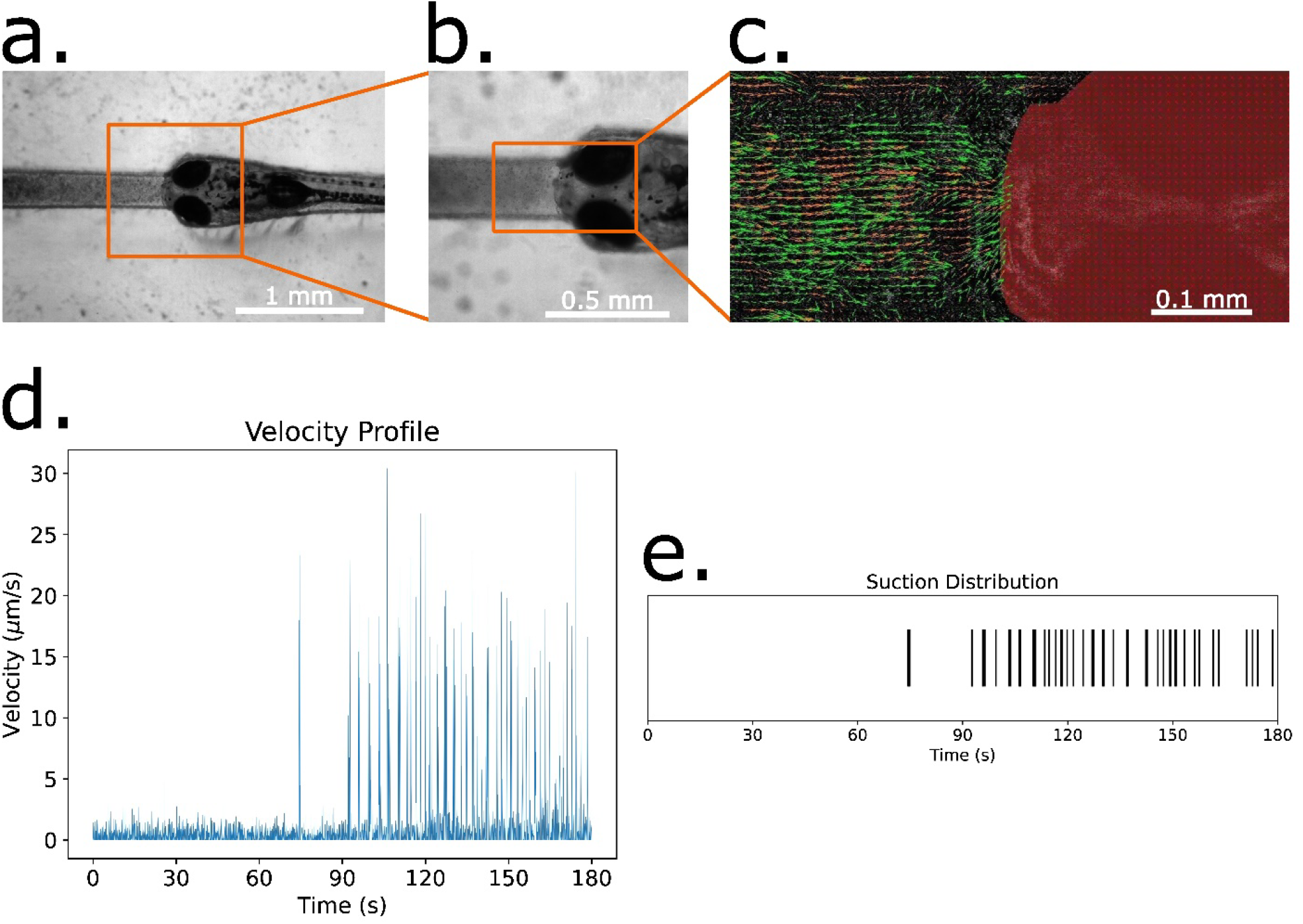
Analysis of food particle suction and velocity in zebrafish larvae. (a) Zebrafish larva immobilized within a microfluidic slot designed using an agarose stamping device for food-suction analysis. (b) Magnified view of the larva within the slot, highlighting its constrained position for controlled observations. (c) Particle Image Velocimetry (PIV) analysis of food particle motion near the larva’s mouth. Velocity vectors represent particle displacement within a single frame. (d) Temporal profile of the average velocity of food particles across all frames during a 3-minute experiment. (e) Suction event distribution over time, extracted from velocity measurements by identifying peak velocity occurrences as suction events.

To quantify food particle motion during suction events, we used an upright epifluorescence microscope setup (Olympus Corporation, Japan) with a 10× air objective (Olympus Corporation, Japan) and a high-speed sCMOS camera recording at 33 frames per second (fps), controlled by HC Image Live software (version 4.5.0.0, Hamamatsu Corp., Japan). The feeding activity of the larva was recorded for 3 minutes, and the video was subsequently converted into individual frames using ImageJ. We then applied Particle Image Velocimetry (PIV) analysis using the PIVlab toolbox in MATLAB to extract food particle velocities in each frame (Figure 5c).

A 200 × 200 μm2 region of interest (ROI) was defined directly in front of the larva’s mouth, and the average velocity of all food particles within the ROI was calculated for each frame. This value was assigned as the frame velocity. Using Python, we converted the frame-based velocity data into a continuous temporal velocity profile over the 3-minute recording (Figure 5d). The velocity profile for this larva revealed a noticeable increase in suction events in the latter half of the experiment, indicated by sharp velocity peaks. To identify suction events, we extracted velocity peaks from the velocity profile and validated them manually by reviewing the recording in ImageJ. The final suction event distribution over time is shown in Figure 5e, providing insights into the feeding behavior and suction frequency of the zebrafish larva.

### Reusability of Stamps and Devices

A benefit of this method is that both the stamps that are developed and the Agarose Stamped Devices (ASD) are reusable. The stamp can be used to make multiple ASD’s and the ASD’s can be used for multiple trials depending on the experiment. If measures are taken to keep the ASD from losing any moisture (ie: enclosing the petri dish or keeping it submerged in water) the ASD will retain the dimensions for its features for several weeks.

### Ease of Use of Devices

Placing a larva into the Agarose Stamped Device (ASD) can take a matter of seconds. Depending on the features chosen, a carefully placed larva via pipetting could be directly deposited onto the larva holding compartment (LHC) of the device. Still, larva placed slightly outside the LHC can simply be aligned and moved into the LHC with a simply made hair loop device (HLD).

Some caution must be taken when utilizing a device with high throughput features as it can increase difficulty in placing larvae. LHC features placed near each other (<10 mm) requires care that no excess water is provided when the larva is pipetted into the ASD as the excess water could displace other larvae from their LHC.

Alternate strategies to utilize high throughput LHC features:

- Reduce density of LHC Features
- Place all larvae for the trial into the ASD before moving and aligning them into the LHC features

